# Assessing the influence of edge effects on macrofaunal contributions to decomposition rates across forest-field ecotones

**DOI:** 10.64898/2026.06.24.734239

**Authors:** Trisha E. Niles, Carmelia Taheri, Robert W. Buchkowski

## Abstract

Understanding the relationships between soil macrofauna and decomposition is crucial for predicting how land-use change impacts ecosystem function in fragmented systems. This is because soil macrofauna affect decomposition and also respond to the changes in abiotic conditions across habitat gradients. This study investigates edge effects on the macrofauna contributions to decomposition across forest-field ecotones. We used bait lamina assay to quantify aboveground and belowground feeding activity of soil macrofauna in Autumn 2025 in three deciduous forest-old field ecotones and one coniferous forest-old field ecotone, in Southwestern Ontario, Canada. Vegetation diversity and composition, LAI and soil characteristics (i.e., soil organic matter, pH, temperature and moisture) were measured at each plot along the ecotone. Pitfall trap data collected in Summer 2025 at the same sites were used to characterize macrofauna communities. We used generalized linear mixed effects models to estimate the effect of distance to edge, site, and depth into the soil on bait lamina consumption and soil macrofauna, with transect nested within site as random effects. Consumption activity increased with distance into the forest from the field, with the edge representing an intermediate; and, decreased with increasing depth into the soil. In contrast, soil macrofauna abundance, especially isopods, decrease with distance into the forest from the field. These trends varied significantly across sites, so that consumption activity and abundance sometimes remained constant across the ecotone (i.e., site × distance interaction). The results demonstrate that macrofaunal contributions to bait consumption varied along the ecotone, shaped by interacting environmental gradients and shifts in community composition unique to each site.

## 1. INTRODUCTION

Decomposition is a fundamental ecosystem process whose rate varies across habitat gradients (Swift et al. 1979; Lavelle et al. 1993; Wardle 2002; Coleman et al. 2017). This variation may cause differences in ecosystem functioning and services (Mitchell et al. 2014). Modern ecology recognizes these habitat gradients as *ecotones*, a term first coined by Clements (1905) that has since evolved to describe areas where communities, ecosystems and ecoregions experience transition (Kark 2012).

Differences in ecosystem properties seen at an ecotone—known as *edge effects*—alter decomposition. Such changes include microclimate, litter quality and soil fauna communities, which we expect to be different at ecotones (Swift et al. 1979; Gessner et al. 2010; Remy et al. 2017). For example, Didham and Lawton (1999) found that litter moisture and air temperature increased, while litter depth decreased, when moving from the forest edge into interior habitat. Edge effects were found to be two to five times greater at open field edges than edges that were closely buffered by another forest. Furthermore, Davies-Colley et al. (2000)’s study of a native broadleaf rainforest-grazed pasture ecotone in New Zealand, found greater foliage density at the edges, and that light exposure and soil temperature had more variable gradients diurnally at points at least 40 metres into the forest. Gehlhausen et al. (2000) found similarly in their study of two isolated forest fragments in east-central Illinois that humidity, soil moisture and light availability changed with distance to edge and aspect (i.e., compass directionality of slope). Additionally, Riutta et al. (2015) found soil moisture decreased from the forest interior towards the edge. Therefore, we expect ecotones to produce unique patterns in the abiotic drivers of decomposition and macrofauna community composition (Anderson 1975, 1977).

Heterotrophic soil organisms are large contributors to decomposition (Wardle, 2002). Since their density changes across environmental gradients, this implies that ecosystem services they provide also change. The decomposer community is composed of soil fauna existing at several orders, in accordance to their size and functional group: microfauna (<0.1 mm, such as *Nemotoda* (nematodes) and protozoa); mesofauna (0.1-2.0 mm, such as *Acari* (mites) and *Collembola* (springtails)); and, macrofauna (>2.0 mm, such as *Oligochaeta* (earthworms), *Isopoda* (isopods), and *Diplopoda* (millipedes); Swift et al. 1979, Wardle 2002). Macrofauna operate at larger temporal and spatial scales than microfauna and mesofauna; possess the ability to transport matter directly to soil substrates; produce additional substrates and food resources for other soil organisms; and, affect microbial communities by grazing (Anderson 1975; Lavelle et al. 1993).

The presence of soil macrofauna exerts a large influence on decomposition due to their litter shredding and geoengineering effects, which implies that they might be an important driver of changes in decomposition across ecotones (Swift et al. 1979; Wardle 2002; Bétard 2021). Vasconcelos and Laurance (2005) tested if leaf litter decomposition rates were affected by changes in microclimate, plant composition and soil fauna by comparing mass loss from two types of leaf mixtures from interior primary and second-growth forests, as well as macroinvertebrate incidence data. They found in their faunal exclusion experiment that variations in plant litter nitrogen to carbon ratios and lignin content as well as the exclusion of soil invertebrates resulted in slower decomposition rates by up to 5% in successional forests and 35% in old-growth forests. They concluded that resource quality determines the range of decomposer community that forms over spatial scales and the processes they oversee, based on the structure of and microenvironment around detrital particles, which are factors that have been established to be influenced by the edge effect. Remy et al. (2017) explored edge effects on nutrient release (of nitrogen, phosphorus and carbon to nitrogen ratio) and litter decomposition in four Belgian temperate forest stands using interior-to-edge transects. Using litterbag experiments with litter predominantly from oak and pine stands, they found litter at the edge lost 87% and 37% more mass, respectively, than litter in interior habitat.

The dynamism of microclimate at habitat boundaries (as a result of edge effects) can alter resource availability for edge-inhabiting organisms, thereby influencing soil fauna abundance and diversity. These findings underscore the potential for ecotone abiotic conditions to alter soil macrofaunal contributions to decomposition; yet, compared to microbial decomposition, this topic remains relatively underexplored. With limited studies on the role of soil macrofauna across forest-field ecotones, and studies calculating decomposition rates for entire ecosystems based on measurements from interior habitat only, this limits our ability to predict decomposition dynamics in fragmented systems. Given this context, this study asked how edge effects alter contributions of macrofauna to decomposition rates across forest-field ecotones. We hypothesized that their contributions to decomposition would be greatest in the forest habitat interior and lowest at habitat edges, given higher litter quality and relatively more stable microclimatic conditions to support diverse macrofaunal communities in the forest. If this hypothesis is correct, we expected that macrofauna would consume more added resources (i.e., bait) in forest interior habitat plots than edge plots along the ecotone-spanning transects.

## 2. METHODS

### 2.1. Study Sites

This study used four ecotones, each containing an old field and adjacent forest. Baldwin Flats adjacent to Gibbons Park in London, ON (*43.001484, -81.270353*) features an old field connected to a temperate deciduous forest. FRAM Lands, also in London (*43.005526, - 81.284170*), is an expansive old field bordered by temperate deciduous forests. Elginfield Observatory (*43.191939, -81.315897*) features an old field habitat sharing a boundary with an evergreen coniferous forest on one side and a temperate deciduous forest on another. To distinguish the two ecotones in this one site, we will refer to the former as Elginfield Coniferous, and the latter as Elginfield Deciduous. These field sites were selected on the basis that they are owned by the university, easily accessible, and large enough to accommodate the chosen experimental design. This creates a study environment representative of urban and peri-urban natural areas.

### 2.2. Experimental Setup

Data collected for this study come from two experimental periods with similar experimental setups. In the summer of 2025, we collected data on the abundance and richness of resident macrofauna across these four ecotones to understand the macrofauna community at each site using pitfall traps. Transects were established in mid-June 2025, and data collection began in late June-early July. Two 80-m transects were set up at each of the four ecotones. Transects were established 20 m away from each other perpendicular to the forest-field edge, spanning 30 m into the forest and 50 m into the field. This was achieved by using a Haglöf Vertex 5 (ver. 2020) hypsometer to measure distance and a compass to set the azimuth. If visualized as a cartesian plane, the midway point of the line formed is the origin (i.e., 0 m) at the forest-field edge. Six 1-m^2^ quadrats were established along the transect line at the following positions: at the edge (i.e., 0 m), 10 and 30 m into the forest interior; and, 10, 30 and 50 m into the field interior.

In the fall of 2025, we performed a bait lamina assay and surveyed vegetation/soil microclimatic variables in the same four ecotones. Old transects were dismantled and two new transects were established on September 29^th^/30^th^, 2025 to avoid areas where macrofauna were removed by our pitfall traps. Data collection began once bait lamina strips were deployed in mid to late October. We used a similar design as before, except the bait lamina set-up had transects spanning 25 m into each ecosystem interior rather than 50 m, with each having five plots at the following positions: at the edge (0 m), 10 and 25 m into the forest interior; and, 10 and 25 m into the field interior (see Figure 1). Across both studies, we avoided placing plots directly on or near walking trails and bike paths, since introduced disturbance and contaminants can reduce or eliminate soil animal feeding activity and abundance (especially of the *Oligochaeta, Diplopoda, and Mollusca* (molluscs) groups; Vorobeichik and Bergman 2020b).

**Figure 1.**
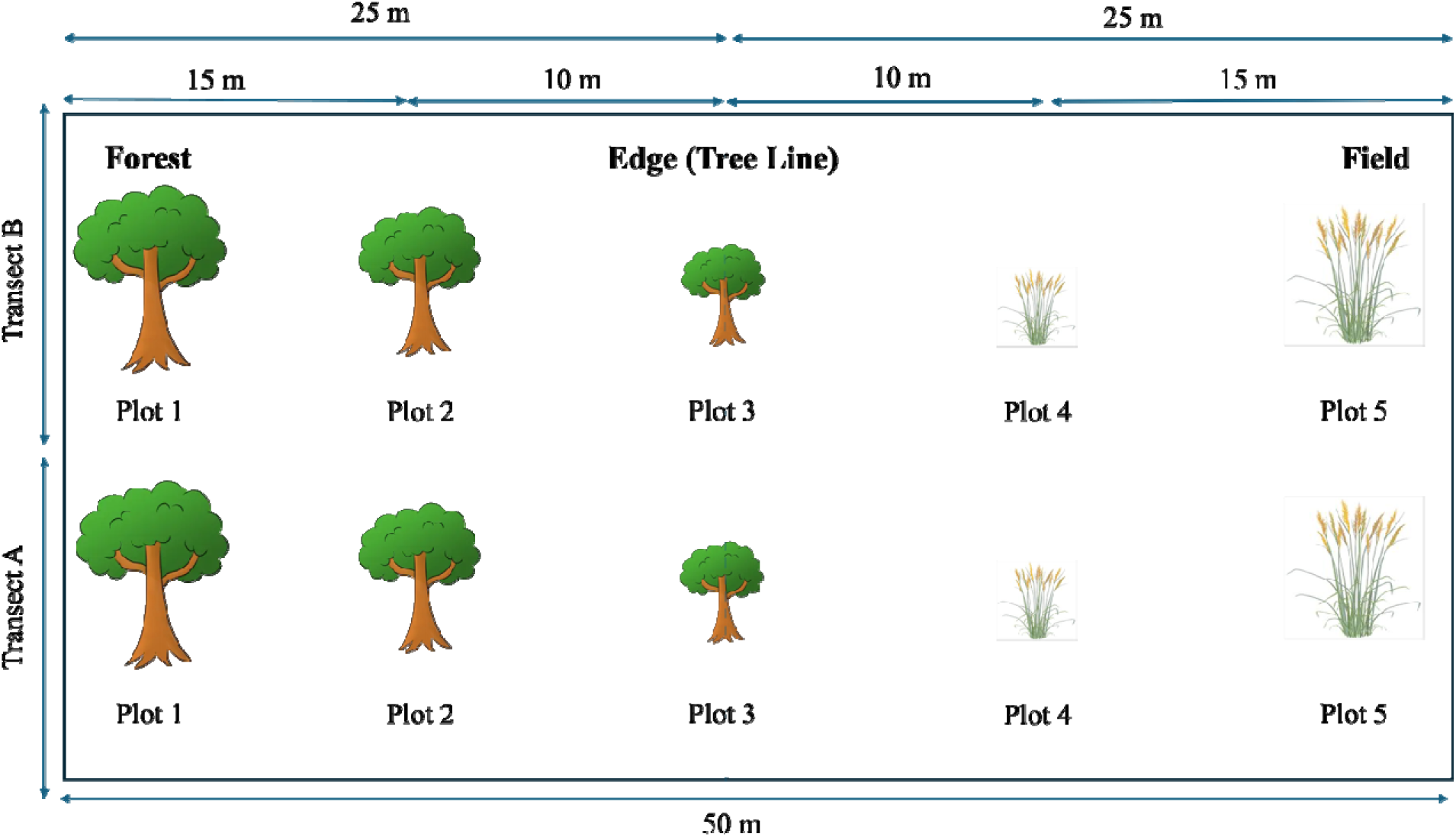
Schematic of experimental design within a site, feature of 5 plots arranged at various distances relative to the tree line (i.e., forest-field edge). During the soil macrofauna survey, an additional plot was added 50-m into the field and plots 1 and 5 were 30-m from the edge instead of 25-m.

### 2.3 Data Collection

#### 2.3.1. Bait Lamina Assay

We quantified decomposer activity using the bait lamina (BL) test. The loss of bait material (i.e., area consumed) from the strip apertures, as well as distribution of consumption vertically along each strip, were assessed and serve as a proxy for decomposition rate (von Törne 1990; ISO 2016). The BL strips are made of red PVC material, sized 120 mm X 6 mm X 1 mm, with 16 apertures around 1.5 mm in diameter spaced 5 mm from each other, and a pointed tip for easy insertion into soil (Podgaiski et al. 2011).

The bait mixture was created using a well-established mixture of cellulose powder, finely grounded wheat bran, and activated carbon, at a ratio of 70:27:3 (Latter and Howson 1977; ISO 2016; Eckert et al. 2025). To make the bait, we weighed components using a calibrated scale, with water added in small amounts while mixing to create a dough-like paste (Kratz 1998; Podgaiski et al. 2011; Riutta et al. 2015). We pressed the bait mixture into the apertures by hand and allowed it to air dry, repeating the process twice, all while checking with a light to ensure no gaps remained unfilled before deployment (von Törne 1990; ISO 2016).

At each plot, we placed three pairs of two BL strips in triangular formation equally distanced from each other (Podgaiski et al. 2011; Riutta et al. 2015); Appendix B, Figure 2). One strip was placed vertically 8 cm deep, with the uppermost aperture at leaf litter-soil level, and was left *in situ* for 14 days. The second strip rested horizontally on the soil surface and was retrieved in 4 days to ensure bait was not entirely consumed before data analysis. We used preliminary trials in Baldwin Flats to determine these deployment times. At the end of the experimental period, we deployed control strips immediately before experimental strip collection to determine baseline loss caused by deployment. Control strips exhibited no significant loss of bait, so the baseline loss is zero for this experiment. We used ImageJ, an image analysis software (Schneider et al. 2012), to quantify the proportion of bait eaten at each aperture from photos taken of the experimental strips.

**Figure 2.**
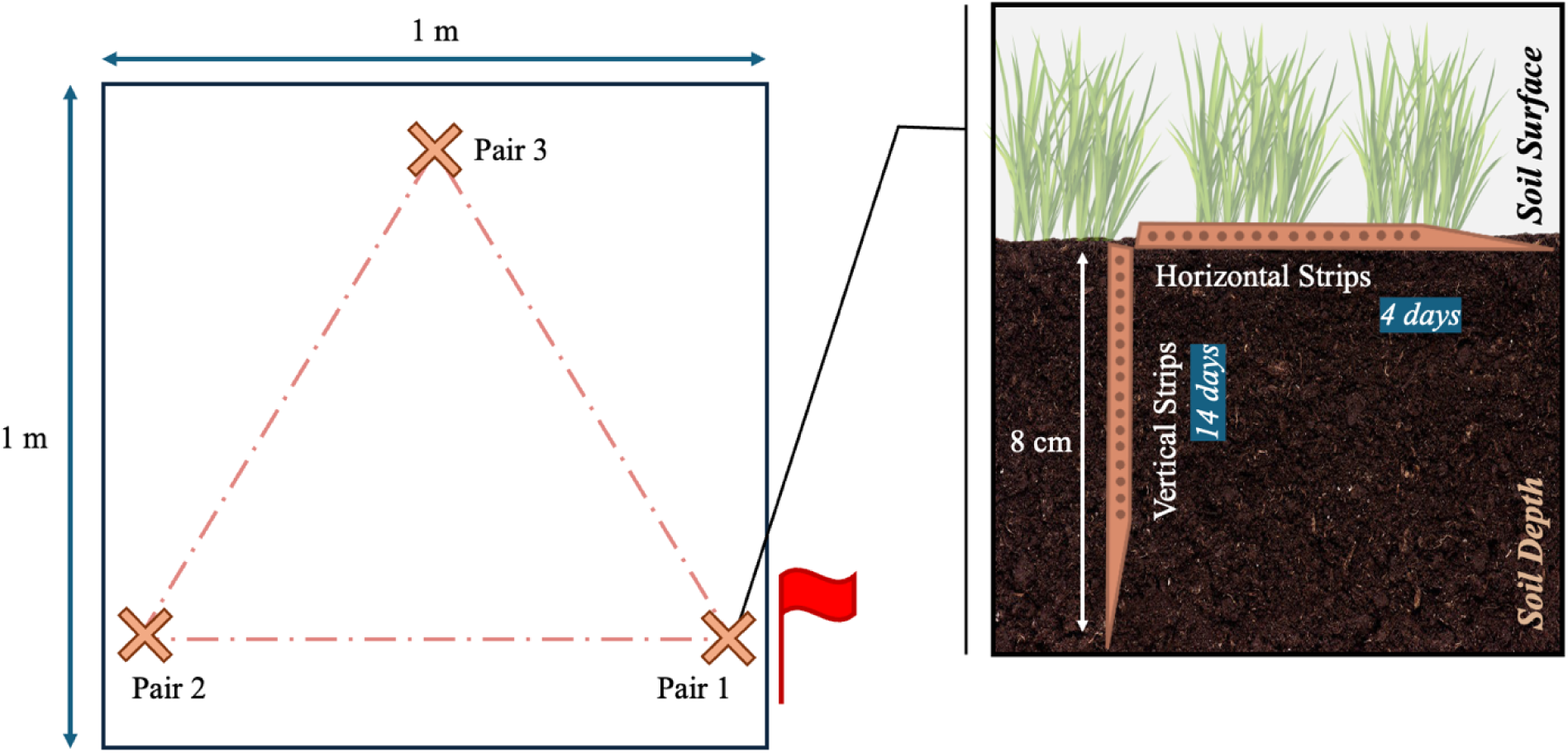
Schematic of bait lamina (BL) setup within each 1 m^2^ plot. BL strips were deployed in pairs: one strip horizontal at the soil surface, and a second strip vertical in the soil 8 cm deep. Three pairs will be setup equidistant to each other in triangular formation within the plot.

#### 2.3.2. Soil Macrofauna Survey

Pitfall traps were set up and active between June 26^th^-July 4^th^, 2025. In the centre of each plot, a 4.7 cm diameter soil core was taken and further dug out to fit the pitfall trap, which had a diameter of 8.4 cm and submerged 12.2 cm deep in the soil so that the upper lip of the trap was level with the soil surface. Once set, we filled the traps with antifreeze to preserve species that were captured. The traps had angled covers made of plastic disposable dinner plates and wooden dowels 10-15 cm above them to prevent rainfall from flooding the traps. The traps were retrieved after 7 days *in situ*. We counted the captured individuals by functional group and then preserved them in Ethanol 95%. Data on all collected animals is available, but only functional groups that likely consumed the bait based on their diet preferences (Potapov et al. 2022) and whose abundance is effectively measured by pitfall traps are presented here.

#### 2.3.3. Soil Temperature, pH, SOM and Moisture

We measured soil organic matter (SOM) by loss-on-ignition using soil collected from cores taken while deploying pitfall traps and passed through a 2-mm sieve (Kalra and Maynard 1991; Hoogsteen et al. 2015, 2018). Soil pH was measured by mixing another subsample of this sieved soil with distilled water and measuring pH using a Thermo Scientific Orion Dual Star pH meter with a combination electrode (Kalra and Maynard 1992; Soil and Plant Analysis Council 1999; Carter and Gregorich 2008).

BL is recommended to be used in temperatures of 5-15 °C (i.e., when soil organisms are active; von Törne 1990; ISO 2016). In addition to collecting point measurement data on soil temperature pre- and post-deployment of BL, we used LogTag HAXO-8 data loggers to record soil temperature every hour beginning November 4^th^ to November 19^th^, 2025, at a depth of around 2.5 cm. We deployed the loggers at each distance along the ecotone in only one of two transects, under the assumption soil temperature fluctuations would be similar along and between the two transects. Since continuous temperature was recorded outside the time period the BL strips were deployed, we reviewed the results to determine *if* the mean and variance of soil temperature changes over the spatial gradient, and how it relates to the point measurements.

Similarly, we collected soil moisture as point measurements using the Hoskin Scientific WET-2 Soil Water Sensor, pre- and post-deployment rather than continuous measurements throughout the duration of the experiment. Thus, it was treated in a similar way as soil temperature, and used as a point of inference only to characterize the variation across the ecotone in this study. Due to the limitations of the data collected, we have only reported site-level averages.

#### 2.3.4. Site Resident Vegetation and Light Exposure

We used the Braun-Blanquet method for floristic surveying as described by Westhoff and van der Maarel (1973) to assess species richness, and percent cover (i.e., the density of individuals of a given species within a plot) as a proxy for species abundance. Species were surveyed by taxa or by family. Leaf Area Index (LAI) at each plot was surveyed using a METER Group LP-80 LAI ceptometer, using open-field above-canopy light exposure as a baseline against light penetration surveyed under all vegetation and leaf litter existing at the soil surface. Both plant and LAI surveys occurred in tandem, between October 11^th^-18^th^, 2025.

### 2.4 Data Modelling and Analysis

#### 2.4.1. Outline of Statistical Methods

Feeding activity (i.e., proportion of bait consumed per strip) is considered the response variable. The explanatory variables are distance to edge, the depth of BL strip aperture into the soil (i.e., soil depth), and site. We also modelled changes in macrofaunal community composition, vegetation composition, measures of soil properties (e.g., SOM, moisture, soil temperature), and LAI at each point along the transects and across sites. This was done to understand how they may change along the ecotone and infer how their gradients may influence BL consumption. All data analysis was completed in _R_ (R Core Team 2025).

Generalized linear mixed effects models (GLMM) were used to account for non-independence inherent in the sampling design, given that BL strip replicates and transects were set up within close proximity of each other in the same plot or site. We used the package *glmmTMB* (McGillycuddy et al. 2025). The *ade4* package was used for multivariate data analysis (Dray et al. 2007), and *vegan* (Oksanen et al. 2001) was used for diversity analysis and ordination. Models were validated and changes in model structure were chosen based on fit statistics using the *DHARMa* package (Hartaig et al. 2024). Summaries and post-hoc tests were performed to assess the significance of relationships between the response and each explanatory variable using standard assumptions of z- and p-statistics, and retrieve coefficients to quantify change in each variable along the ecotone. Data were visually presented using *ggplot2* (Wickham 2016).

#### 2.4.2. Structure of Key Models

Our first model, Model 1, uses an *ordbeta()* distribution and predicts the proportion of bait consumed in each strip using the interaction between strip orientation and site, as well as the mean-centered distance to the field-forest edge. Strip orientation differentiates horizontal and vertical strips. Site was included as a random effect and the number of days of deployment was included as an offset term to correct for the different deployment time for strip orientations.

Our second model, Model 2, analyzes bait consumption in the vertical strips only as a function of the interaction of site and vertical depth into the soil, the interaction of distance to edge and site, and a nested random effect of transect within site. Like Model 1, distance to edge and soil depth were scaled to centre their means. We used a dispersion formula to account for the unequal variance resulting from the interaction of site and soil depth. Likewise, a zero-inflation formula corrected for excess zeroes at certain sites and transects.

Models 3A, 3B and 3C analyze vegetation species richness, abundance and diversity, respectively, as a function of distance to the edge, with a nested random effect of transect within site. Model 3A is fitted to a *poisson*() distribution, while Model 3B and 3C are fitted to a *gaussian()* distribution. Model 4 is a GLMM fitted to a *gaussian()* distribution, measuring LAI as a function of the interaction of distance to the edge and site with a nested random effect of transect within site.

Models 5A and 5B model macrofauna abundance and richness respectively, as a function of distance to the edge, with a nested random effect of transect within site and a dispersion formula to account for unequal variance across sites. Models 5C and 5D use *nbinom2()* or *gamma()* distributions, respectively, to analyze individual species’ abundance as a function of distance to edge and/or site (or the interaction of these two terms), with a nested random effect of transect within site. We used dispersion formulas and/or zero-inflation formulas, if required, to improve model fit.

Model 6 uses a *gaussian()* distribution to SOM as a function of the interaction of site and distance to the edge, with transect as the random effect. A dispersion formula was used to account for the unequal variance resulting from the interaction of site and transect. Model 7 fits soil pH data to a *gaussian()* distribution, as a function of the additive effects of distance and site. We used transect as a random effect, and a dispersion formula to account for unequal variance across sites.

## 3. RESULTS

### 3.1. Bait consumption versus Distance to Edge, Strip Orientation and Site (Model 1)

Average bait consumption across all sites is largest in the forest and gradually deceased with increasing distance into the field (GLMM: z = 2.80, p < 0.01). Consumption at the forest-field edge was found to be an intermediate between forest and field plots. These overall trends were driven by consumption in the horizontal strips rather than consumption in the vertical strips (see Figure 3). There was significantly less consumption in the vertically-placed strips versus horizontally-placed strips at Baldwin Flats after correcting for the different deployment length (GLMM: z = -11.32, p << 0.001). This was relatively consistent across Baldwin Flats, FRAM Lands (GLMM: z = 1.76, p > 0.05), and Elginfield Coniferous (GLMM: z = 1.28, p > 0.10), however, the horizontal strips at Elginfield Deciduous were consumed much more than any of the others (GLMM: -2.32, p < 0.05), leading to a significant interaction between site and strip orientation (see Supporting Information, Table S1).

**Figure 3.**
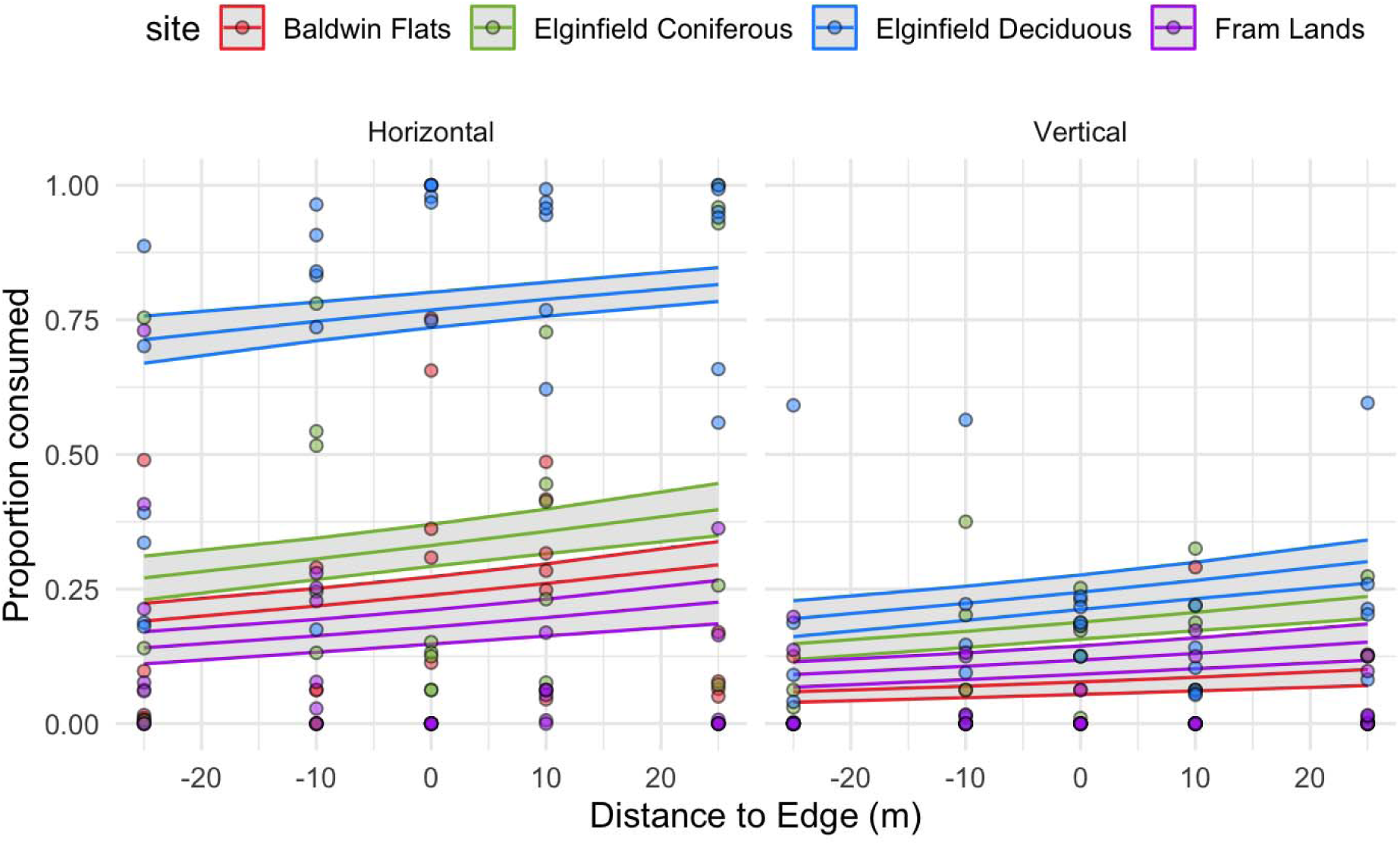
Proportion of bait consumed with distance from the edge (m), in both horizontal (aboveground) and vertical (belowground) BL strips. Negative distance represents increasing distance into the field from the forest edge. Lines show predictions from a generalized linear mixed effects model and points show average consumption at individual plots.

### 3.2. Distance to Edge, Soil Depth and Site versus Bait consumption (Model 2)

Average bait consumption along the vertical strips was greatest within 0-2 cm from the soil surface, gradually deceasing with increasing depth into the soil profile. The results of the model indicate that there is a significant interaction between site and depth in feeding activity in FRAM Lands (GLMM: z = -4.45, p << 0.001), Elginfield Coniferous (z = -2.86, p < 0.01) and Elginfield Deciduous (GLMM: z = -3.55, p < 0.001), driven by the fact that consumption decreased with depth at these three sites; yet it appears that in Baldwin Flats, feeding activity remained constant across soil depth (GLMM: z = 0.181, p > 0.10; Figure 4) Isolating for vertical strip feeding activity, this model also suggests that belowground feeding activity in Baldwin Flats (GLMM: z = -1.98, p < 0.05) decreased and Elginfield Coniferous (GLMM: z = 2.65, p < 0.01) activity increased, with greater distance into the forest from the field, with the edge acting as an intermediate point. Distance to edge was not observed to be a significant predictor of feeding activity in FRAM Lands (GLMM: z = 1.044, p > 0.10), and Elginfield Deciduous (GLMM: z = 1.90, p > 0.05). In all, the model provides evidence for interactions of site with depth, and with distance to edge, in average consumption activity belowground (see Supporting Information, Table S2).

**Figure 4.**
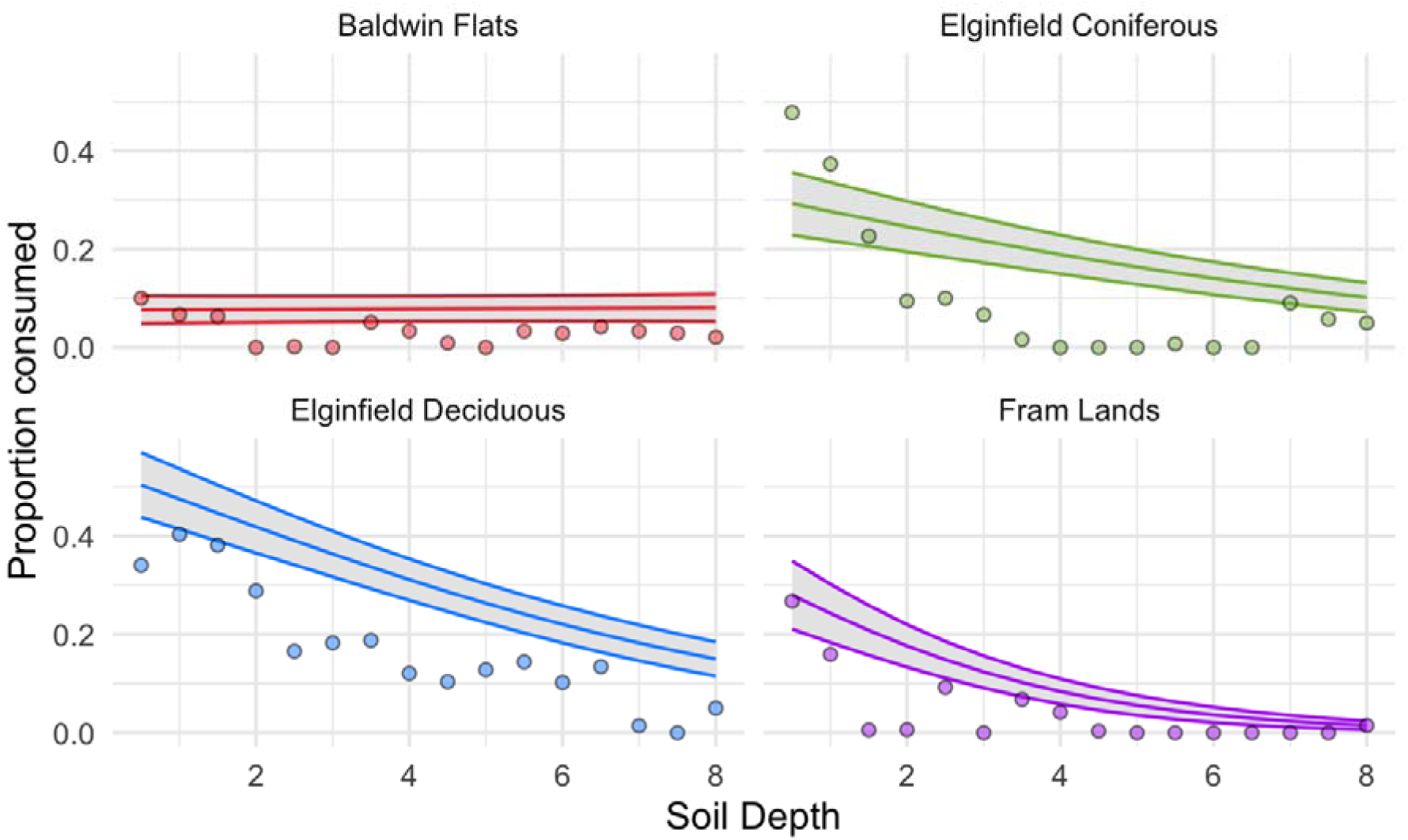
Proportion of bait consumed with soil depth (cm) from vertical BL strips at each site. Lines show predicitions from a generalized linear mixed effects model and points show site-level averages across the entire transect.

### 3.3. Resident Vegetation (Model 3A, 3B, 3C) and LAI (Model 4)

Plant percent cover (i.e., abundance) decreased with increasing distance into the forest across all sites, with the edge acting as an intermediate (GLMM: z = -6.10, p << 0.001; see Supporting Information, Table S3a), as did richness (GLMM: z = -4.31, p << 0.001; see Supporting Information, Table S3b). There was no significant difference in Shannon’s Diversity with distance to edge in Baldwin Flats (GLMM: z = -1.24, p > 0.10) and Elginfield Deciduous (GLMM: z = 0.70, p > 0.10). There is evidence of decreases in Shannon’s Diversity with distance to edge in Elginfield Coniferous (GLMM: z = -2.89, p < 0.01), and in FRAM Lands (GLMM: z = -2.17, p < 0.05; see Supporting Information, Table S3c).

LAI at the soil surface was found to have site-level variations that did not significantly interact with distance to the edge (GLMM: p > 0.05). LAI ranged from 0.90 - 3.71 in Baldwin Flats, 1.82 - 4.85 in FRAM Lands, 1.29 - 4.61 at Elginfield Coniferous, and 1.19 - 3.22 in Elginfield Deciduous. The results of the model (see Supporting Information, Table S4) demonstrate that canopy density when measured from the soil surface remained relatively constant along the gradient; however, it also suggests that site-specific conditions or community structure drive LAI (see Figure 5).

**Figure 5.**
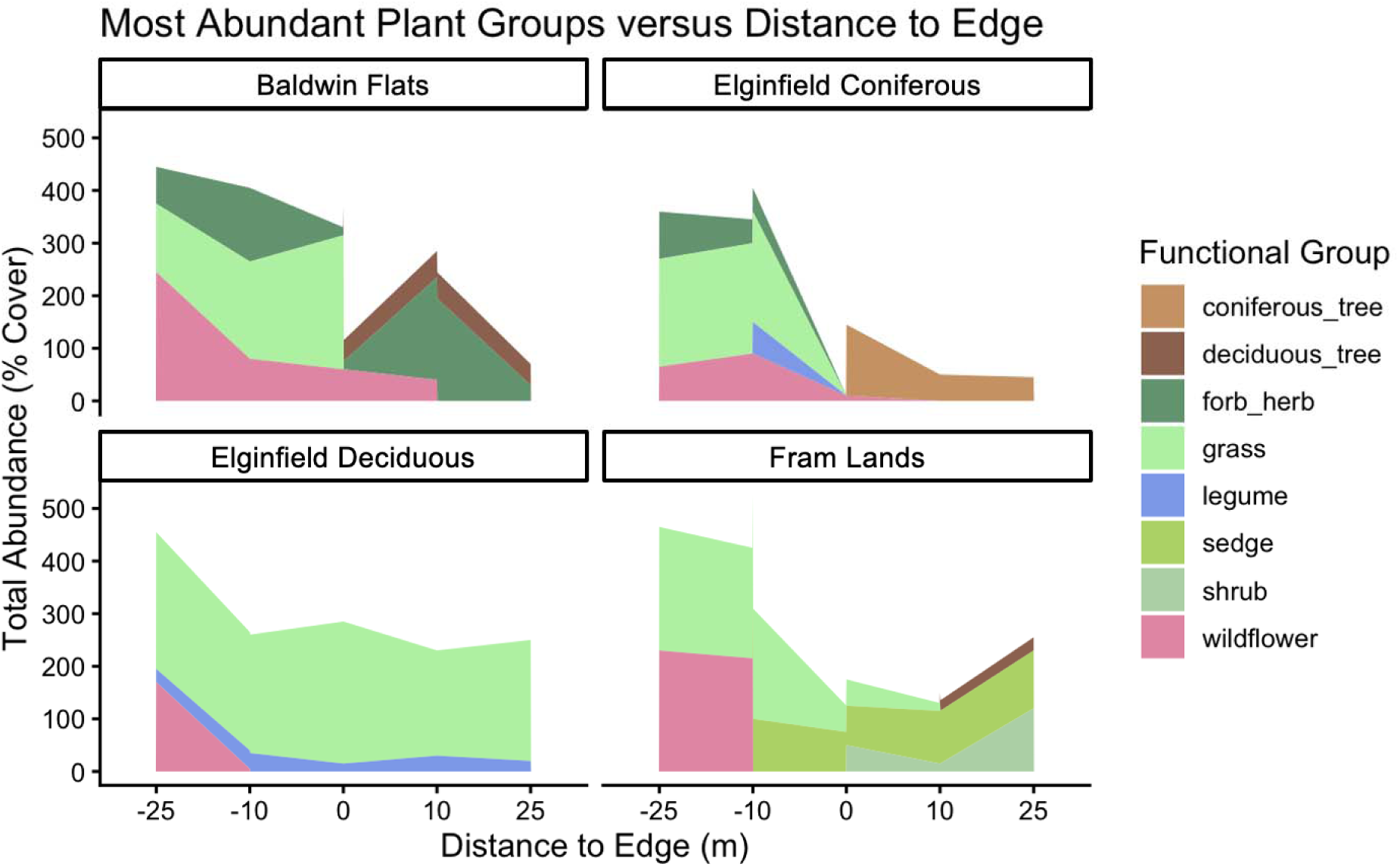
A visualization of the top three dominant vegetation functional groups at each distance to the edge and at each site. Negative distance represents increasing distance into the field from the edge.

### 3.4. Macrofauna Community Composition and Diversity (Models 5A-D)

With increasing distance from the field into the forest, total species abundance (GLMM: z = -2.77, p < 0.006) decreased, with the edge representing an intermediate between the two ecosystems. Species richness appears to follow a similar trend, but not significantly (GLMM: z = -1.887, p > 0.05; see Supporting Information, Table S5a and S5b). Isopod abundance decreased with increasing distance into the forest from the field (GLMM: z = -3.60, p < 0.000313; see Supporting Information, Table S5c), with the same seen with snails (GLMM: z = -6.36, p << 0.01; see Supporting Information Table S5d). Millipedes and slugs showed no significant shifts in abundance with distance from the edge (GLMM: p > 0.05). Over all sites, the invertebrate community composition, as assessed by pitfall traps, changed with distance to the edge driven by changes in the abundance of isopods and snails (see Figure 6).

**Figure 6.**
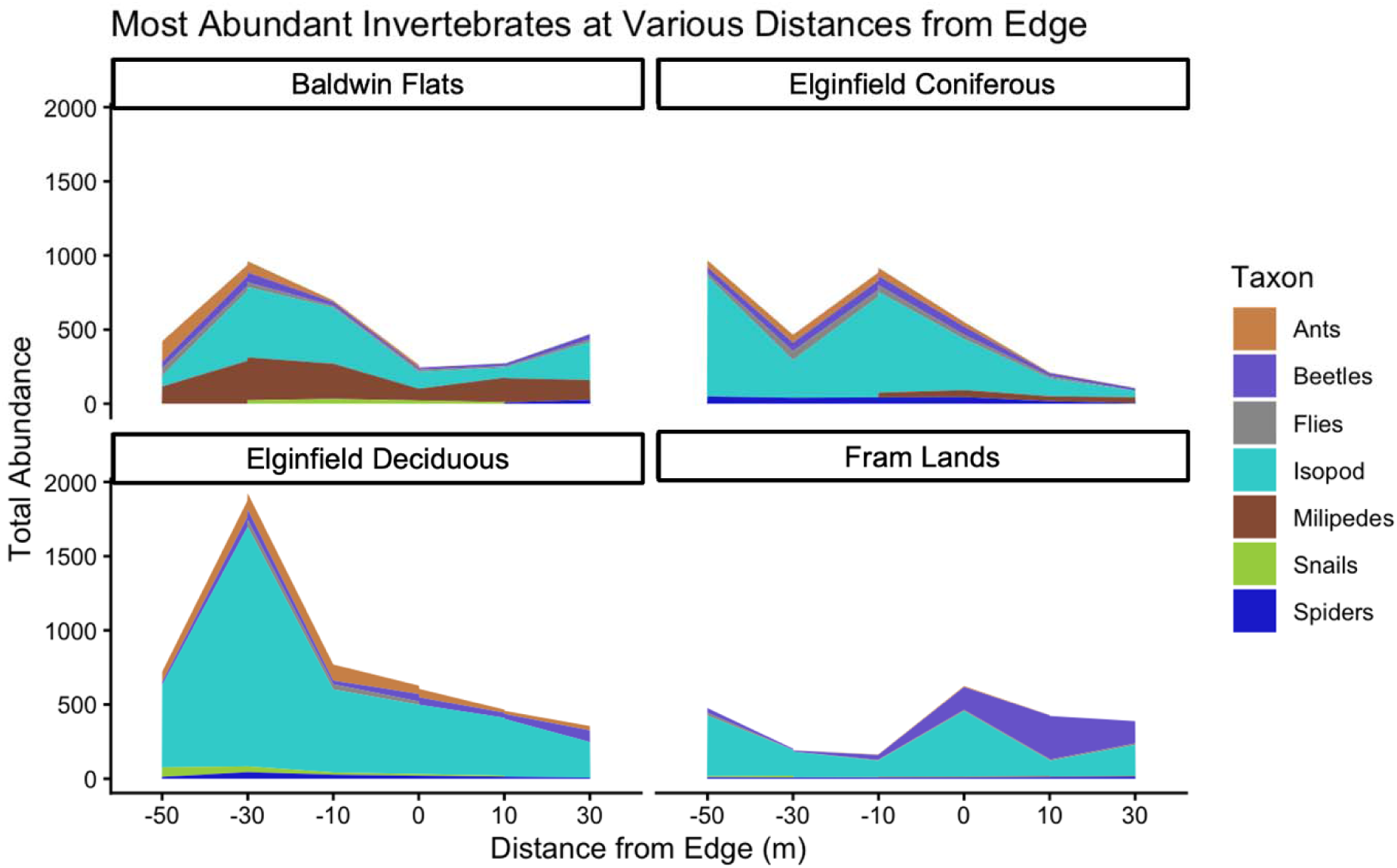
A visualization of the top three dominant macrofauna functional groups at each distance to the edge. Negative distance represents increasing distance into the field from the edge.

### 3.5. Soil Characteristics (Model 6 and 7)

While the experiment was active in October 2025, point measurements taken found that the average soil temperature at each site fell in the following ranges: Baldwin Flats between 10.6°C - 20.1°C; FRAM Lands between 8.9°C - 21.1°C; Elginfield Coniferous between 5.6°C - 13.7°C; and, Elginfield Deciduous between 6.9°C - 17.0 °C. During of the period of time the temperature loggers were deployed in November 2025, the soil temperature at each site fell in the following ranges: Baldwin Flats between 0.9°C - 11.0°C; FRAM Lands between 2.0°C - 9.0°C; Elginfield Coniferous between 0.8°C - 11.2°C; and, Elginfield Deciduous between 2.0°C - 9.4 °C.

Point soil moisture measurements taken at the 4- and 14-day mark of the experiment found that average moisture at each site fell in the following ranges: Baldwin Flats between 2.1% - 20.6%; FRAM Lands between 9.4% - 28.3%; Elginfield Coniferous between 4.0% - 27.9%; and, Elginfield Deciduous between 10.3% - 29.2%.

SOM in Baldwin Flats and FRAM Lands did not significantly vary with distance to the edge (GLMM: z = 1.28, p > 0.05; z = -0.69, p > 0.10). SOM decreased with increasing distance into the forest from the field at the Elginfield Coniferous (GLMM: z = -3.89 p < 0.01), and increased with distance in Elginfield Deciduous (GLMM: z = -3.96, p << 0.001; see Supporting Information, Table S6). This significant interaction between site and distance to the edge meant that the changes in SOM across the ecotone were unique to each site.

The soil pH did not vary with distance to the edge (GLMM: z = -0.80, p > 0.05), but did vary by site: Baldwin Flats had an average soil pH between 7.69 - 7.89; FRAM Lands between 7.10 - 8.10; Elginfield Coniferous between 6.60 - 7.74; and, Elginfield Deciduous between 7.00 - 7.82.

## 4. DISCUSSION

### 4.1. Overall Results

The results partially support our hypothesis that macrofaunal consumption activity was greatest in the forest interior. Contrary to what was expected, the edge had intermediate instead of the lowest bait consumption. Our data reproduced a pattern described by previous studies as an “edge effect”, whereby feeding activity declined from forest interior to the forest edge (Simpson et al. 2012; Riutta et al. 2015). The conflict with our hypothesis was that we observed no subsequent increase in feeding activity from the edge into the field. Consumption of bait above and below the soil surface was found to vary with distance from the edge, soil depth, and site, with site being the most influential. Other bait lamina studies have observed variations in bait consumption at the site-level, attributed to differences in abiotic factors more so than soil faunal community shifts (von Törne 1990, Gongalsky et al. 2003; Filzek et al. 2004; Römbke et al. 2006). In line with current literature, most consumption occurred within the first 0-2 cm from the soil surface (Paulus et al. 1999; Hamel et al. 2007; Riutta et al. 2015; Gongalsky et al. 2003). Overall, our results support existing trends in bait lamina consumption across the forest-field ecotone and soil depth, while significant interactions with site highlight the currently unexplained effect that each unique combination of vegetation, abiotic environment and land use history appears to have on trends in bait consumption.

### 4.2. Vegetation

The forest interior had the highest bait consumption relative to the field, and was dominated by woody species in the overstory, and a greater accumulation of leaf litter (Cadenasso et al. 2003; Loydi et al. 2013). Bait consumption was likely higher in the forest because the thicker litter layer habitat is favourable to the dominant detritivores at our sites (Potapov et al. 2022). The increase we saw may have been exaggerated by measuring activity in Autumn after tree litter fall, when the forest had lots of fresh litter for detritivores and field vegetation had not senesced yet (Carson and Peterson 1990; Facelli and Carson 1991; Loydi et al. 2013). We saw a positive relationship between detritivore abundance and bait consumption across sites, driven largely by the high detritivore abundance at the Elginfield’s deciduous forest site, which contained trees with highly palatable litter, especially in comparison to the Elginfield’s coniferous forest (Cornelissen 1996; De Smedt et al. 2016; Korobushkin et al. 2025). Lastly, routine mowing at our sites produced distinct edges that were a mix of forest and field species rather than any edge-specialists. This may explain why we saw intermediate amounts of consumption activity instead of the decrease in consumption at the edge that we predicted (Facelli and Carson 1991; Cadenasso et al. 2003; Abramoff et al. 2024).

### 4.3. Macrofauna Communities

Average bait consumption was dominated by feeding at the soil surface, which suggests that the most abundant soil surface-dwelling macrofauna at our sites, including isopods and snails, were responsible for the overall trends in bait consumption. Isopods likely dominated bait consumption at our sites given they are particularly proliferous, generalist detritivores that consume leaf litter and woody debris (upwards of 1-4% of deciduous litter pools annually), and are active within the leaf litter stratum (i.e., 0-1 cm depth) of the soil surface (Yang et al. 2020; Potapov et al. 2022). Snails were the second most abundant potential consumer of bait, but given their generalist diet and lower abundance, they likely did not contribute to as large a proportion of bait consumption (Potapov et al. 2022; Schmera et al. 2023).

Both isopods and snails increase in abundance from the forest into field, which was opposite the trend that we observed in bait consumption. This appears unexpected because a larger isopod population should have led to greater average consumption activity (Riutta et al. 2012; De Smedt et al. 2016). However, interpreting this as an unexpected change in their per capita feeding activity may be premature because the population surveys and bait consumption measurements were taken during different seasons and detritivore diversity and abundance fluctuates with season even if dominant taxa remain consistent (Trifonovayr et al. 2015; D’Hervilly et al. 2021). So, our results suggest that either forest isopods were eating more surface bait per capita than those in the field; or systematic biases, such as unrepresentative sampling by pitfall traps across the gradient or population size changes between seasons, created an artificial trend.

Flies, beetles, spiders and crickets are mostly recognized as omnivorous and/or carnivorous, occupy many trophic levels (Potapov et al. 2022), and were relatively rare compared to isopods, so they likely did not consume large amounts of our bait. Earthworms would also be involved in bait consumption (Zimmer et al. 2005; Gongalsky et al. 2008; Vorobeichik and Bergman 2020b), millipedes and slugs likewise (Golovatch and Kime 2009; Potapov et al., 2022). Their abundances may have not been captured well using pitfall traps in our study (Mesibov et al. 1995; Druce et al. 2004), so we cannot properly couple them to observed feeding activity changes with distance to edge.

Additionally, the lowest consumption belowground was observed at Baldwin Flats, the site with the greatest human use, likely a result of disturbances inherent to its land use history as an old farm and current use as a high-traffic urban park. Maintenance, vegetation/soil surface trampling by biking and walking paths, and littering introduce contaminants to the soil system, of which has generally been observed to push soil fauna to exclusively occupy the litter layer. This leads to weak decreases in feeding activity with soil depth compared to relatively undisturbed or unused areas (Vorobeichik and Bergman 2020a), and the creation of an unhospitable environment that reduces abundance of macrofauna (Filzek et al. 2004; Vorobeichik et al. 2019).

### 4.4. Soil Characteristics

We found no robust trends in soil moisture, temperature and pH with distance to the edge, which contradicts observations from past literature (Didham and Lawton 1999; Davies-Colley et al. 2000; Gehlhaussen et al. 2000; Riutta et al. 2015). The present study cannot fully assess the role of these factors given the limited data. However, our moisture and temperature results may be an effect of season, where cool autumn conditions made the forest and field more similar. In general, soil temperature and moisture interact with soil depth to influence soil fauna activity.

Greater activity at the soil surface is expected in the autumn, as the higher moisture and lower temperature conditions are favourable for macrofauna by limiting desiccation (Rożen et al. 2010). Past BL studies found consumption varied with space based on soil temperature and moisture, while the depth of consumption varied with soil texture and compaction (Gongalsky et al. 2003; 2008). While not observed in the present study, literature describes soil pH variations to be a function of distance to edge and site, reflecting differences in vegetation and thus litter quality and type changes with turnover (Filzek et al. 2004; Abramoff et al. 2024). However, soil temperature and pH were not observed to vary with space and depth as consistently as soil moisture in past studies (Simpson et al. 2012; Riutta et al. 2015). Since our study found significant variation across sites and our results conflict somewhat with the predictions made by previous studies—albeit those that did not measure trends in bait consumption all the way into adjacent fields. Further exploration of the edge effect on soil macrofauna, decomposition, and SOM dynamics will be crucial to understand these fragmented, urban ecosystems (Filser et al. 2016; Frouz 2017).

### 4.5. Limitations

Several limitations should be considered when interpreting our results. This project was spatially and time-constrained to four ecotones within three sites and one observation period; thus, data was representative of a fragment of the larger ecoregion. Furthermore, BL feeding cannot be fully attributed to any specific macrofauna or macrofaunal group (Vorobeichik and Bergman 2023). Additionally, macrofaunal gamma-diversity may not vary as much seasonally, yet faunal abundance fluctuates with season such that it may be higher in autumn than summer due to changes in precipitation patterns (Riutta et al. 2012; Ball et al. 2025). This introduces an issue when using the macrofaunal abundance in the summer to explain patterns of BL consumption in the autumn. There is also the inability of pitfall traps to capture the abundance of groups below the leaf litter layer or less mobile soil animals (Mesibov et al. 1995). Lastly, we only took point measurements of soil temperature and moisture during the BL deployment period, and found them to be highly variable. Soil temperature loggers deployed afterwards provided limited information about differences within and between sites. Thus, we could not carry out an analysis of these key soil characteristics and this may have exaggerated the effect of site that would otherwise have been explained by soil temperature and moisture.

### 4.6. Conclusion

This study demonstrated that macrofaunal contributions to decomposition vary along forest-field ecotones, and that the strength of this trend was unique to each site. This highlights the importance of considering site-level trends in detritivore populations when making management plans, especially when maintaining their contribution to decomposition and nutrient cycling is a priority. A future study could explore the contributions of particular soil macrofaunal groups through triangulating both indirect and direct measurements of activity and sampling from more sites over several time periods to better capture the temporal and spatial changes in soil fauna feeding activity in fragmented urban ecosystems.

## ACKNOWLEDGEMENTS

We thank members of our laboratory group, Western University’s Nutrient Modelling & Movement Lab, for their tireless support throughout the experimental set-up and data collection phases. Particularly, R. Nayudu, H. Osmani, S. Zaman, T.J. Chakma, E. Gillis and P. Ferguson for help in the field. We also thank Z. Lindo, H. Henry and B. Souriol for providing their input on the project’s planning and progression. Funding for this project was provided by grants to R.W.B. from the Natural Science and Engineering Research Council of Canada and the Canada Research Chairs program (Grant Numbers: RGPIN-2024-05238 & CRC-2023-00334).

## DATA AVAILABILITY

The data that support the findings of this study are publicly available and accessible from 10.6084/m9.figshare.32804324.

