## Supporting Information for "Assessing the influence of edge effects on macrofaunal contributions to decomposition rates across forest-field ecotones"

**Supporting Information – Main Statistical Model Results**

**Table S1.** Results of a generalized linear mixed-effects model examining the effects of the interactions between strip orientation, site, and the mean-centered distance to the field-forest edge on bait consumption. Site was used as a random effect. BL = bait lamina; EO = Elginfield Observatory (Con. = Coniferous; Dec. = Deciduous); FL = FRAM Lands. The intercept is horizontal BL strip orientation at Baldwin Flats.

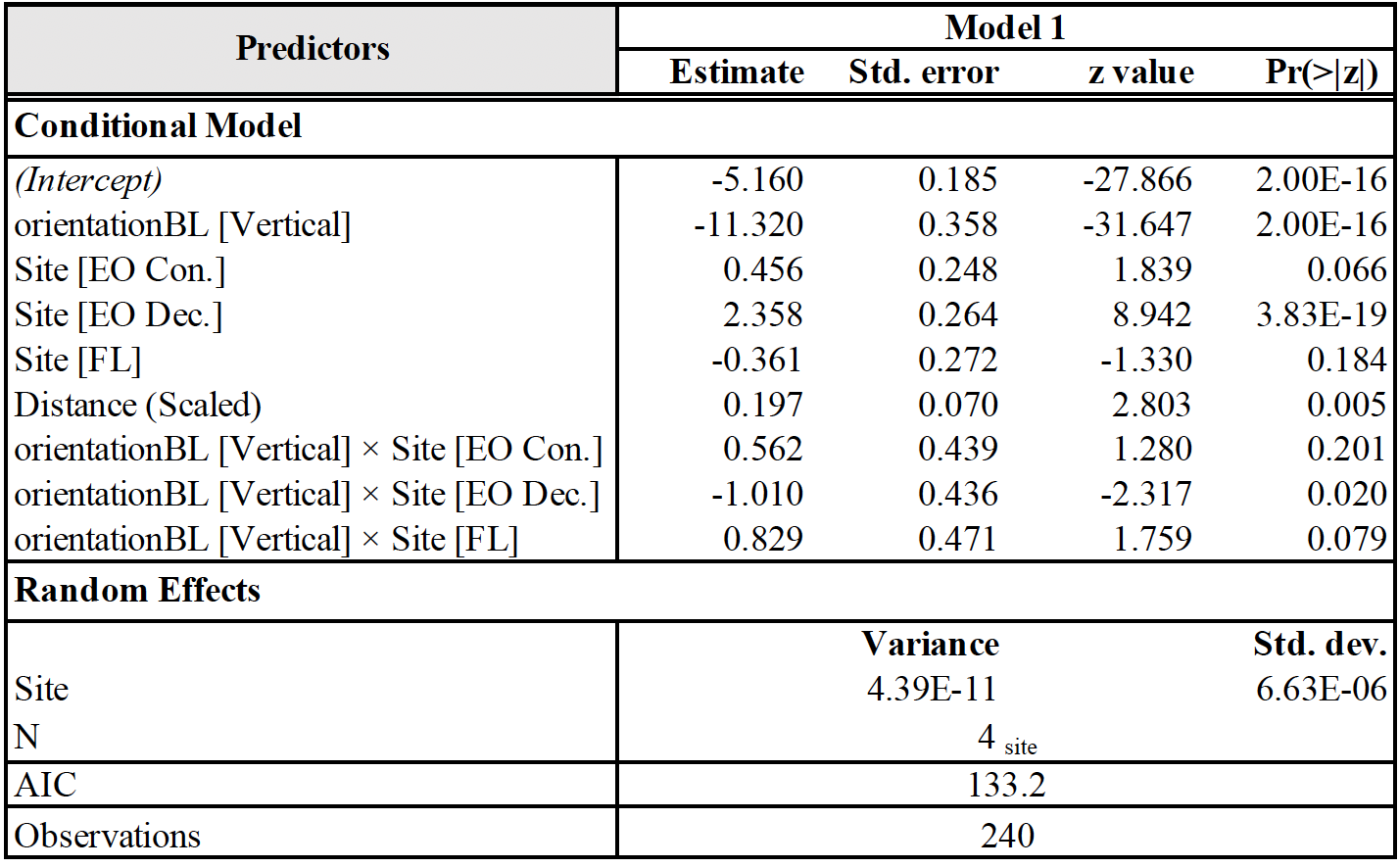

**Table S2.** Results of a generalized linear mixed-effects model examining the effects of interactions of site, mean-centered soil depth, and the mean-centered distance to the field-forest edge on bait consumption. Site and transect nested within site were used as random effects. BL = bait lamina; EO = Elginfield Observatory (Con. = Coniferous; Dec. = Deciduous); FL = FRAM Lands. The conditional model intercept is the baseline response of Baldwin Flats assuming scaled distance to edge and scaled depth equal zero.

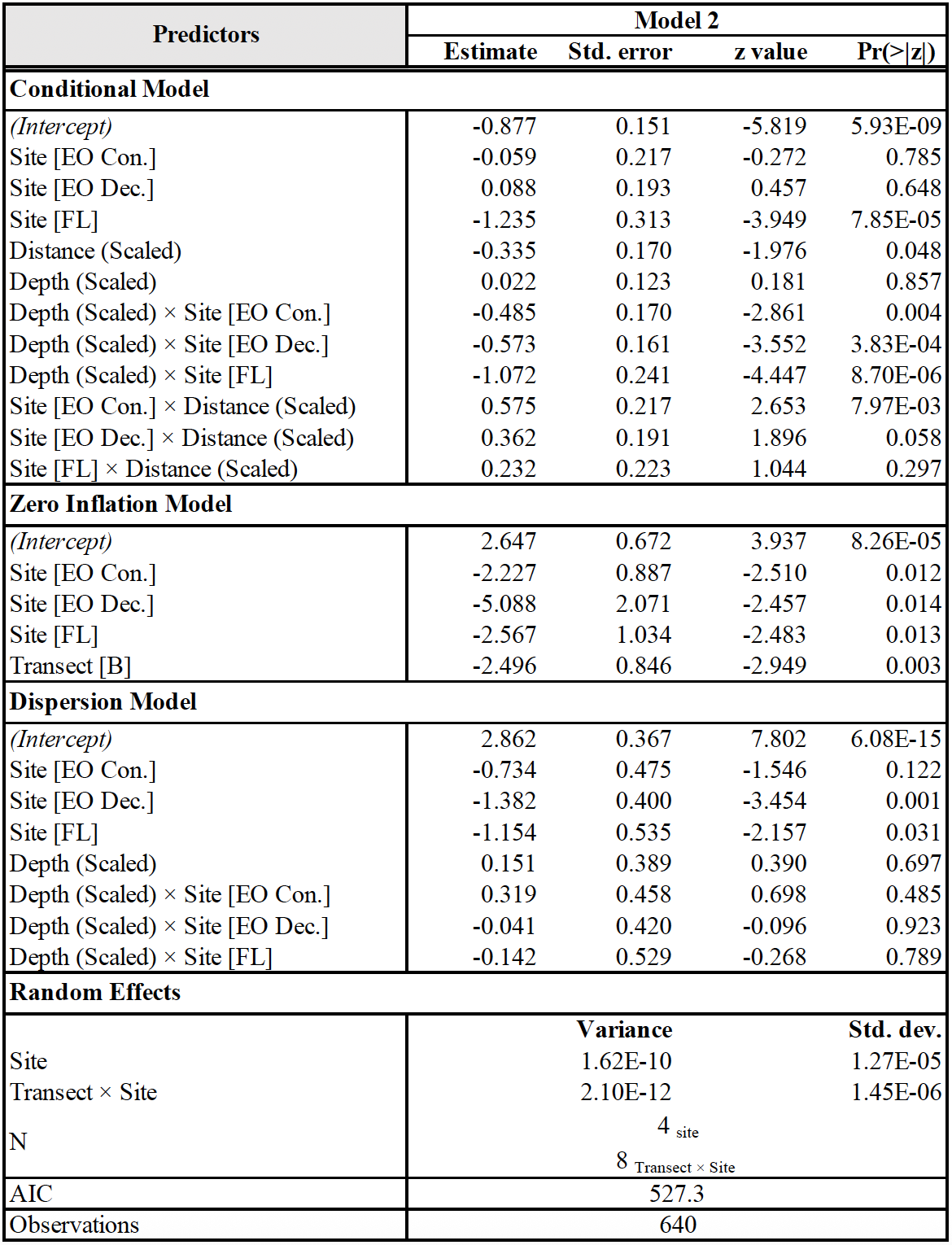

**Table S3a**. Results of a generalized linear mixed-effects model examining the effects of site and distance to the field-forest edge on vegetation abundance (i.e., % cover). Site and transect nested within site were used as random effects. The intercept is the estimated abundance at the edge.

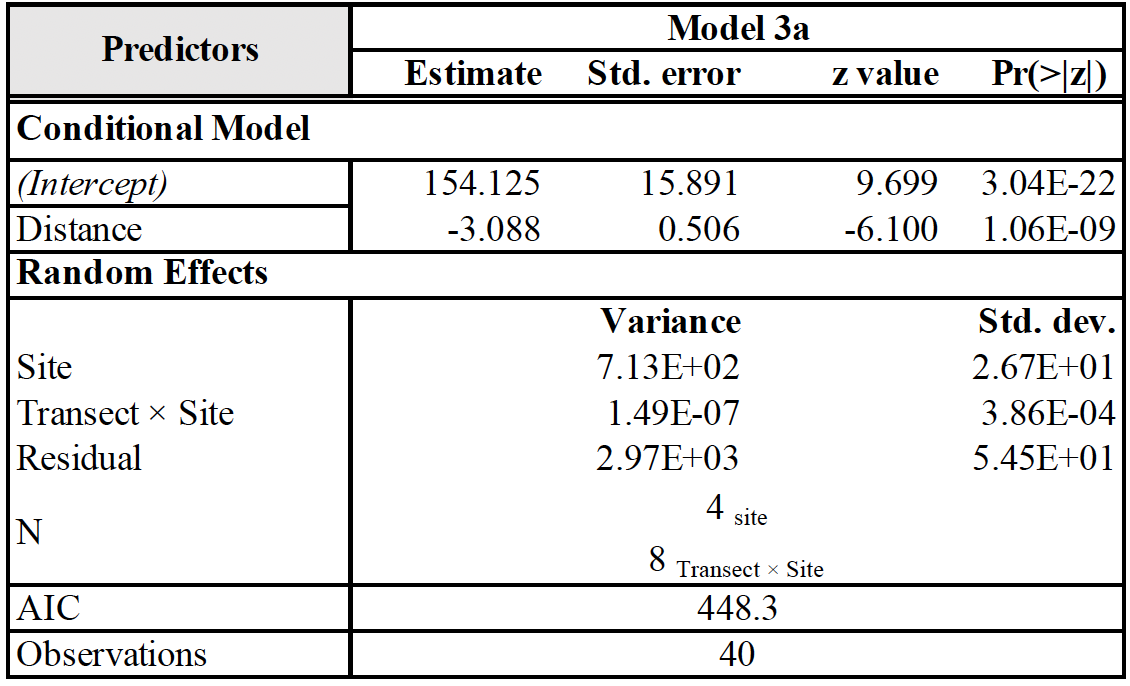

**Table S3b.** Results of a generalized linear mixed-effects model examining the effects of site and distance to the field-forest edge on vegetation abundance. Site and transect nested within site were used as random effects. The intercept is the estimated richness at the edge.

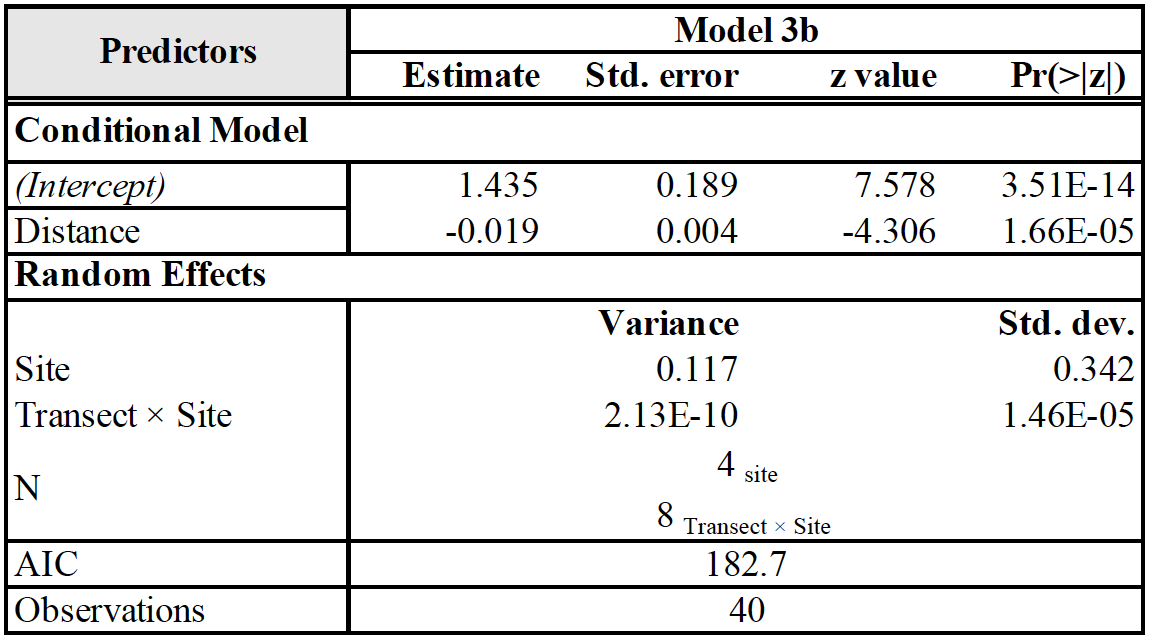

**Table S3c.** Results of a generalized linear mixed-effects model examining the effects of the interaction of site and distance to the field-forest edge on vegetation Shannon’s diversity. Site and transect nested within site were used as random effects. EO = Elginfield Observatory (Con. = Coniferous; Dec. = Deciduous); FL = FRAM Lands. The intercept is the Shannon’s Diversity at Baldwin Flats’ edge.

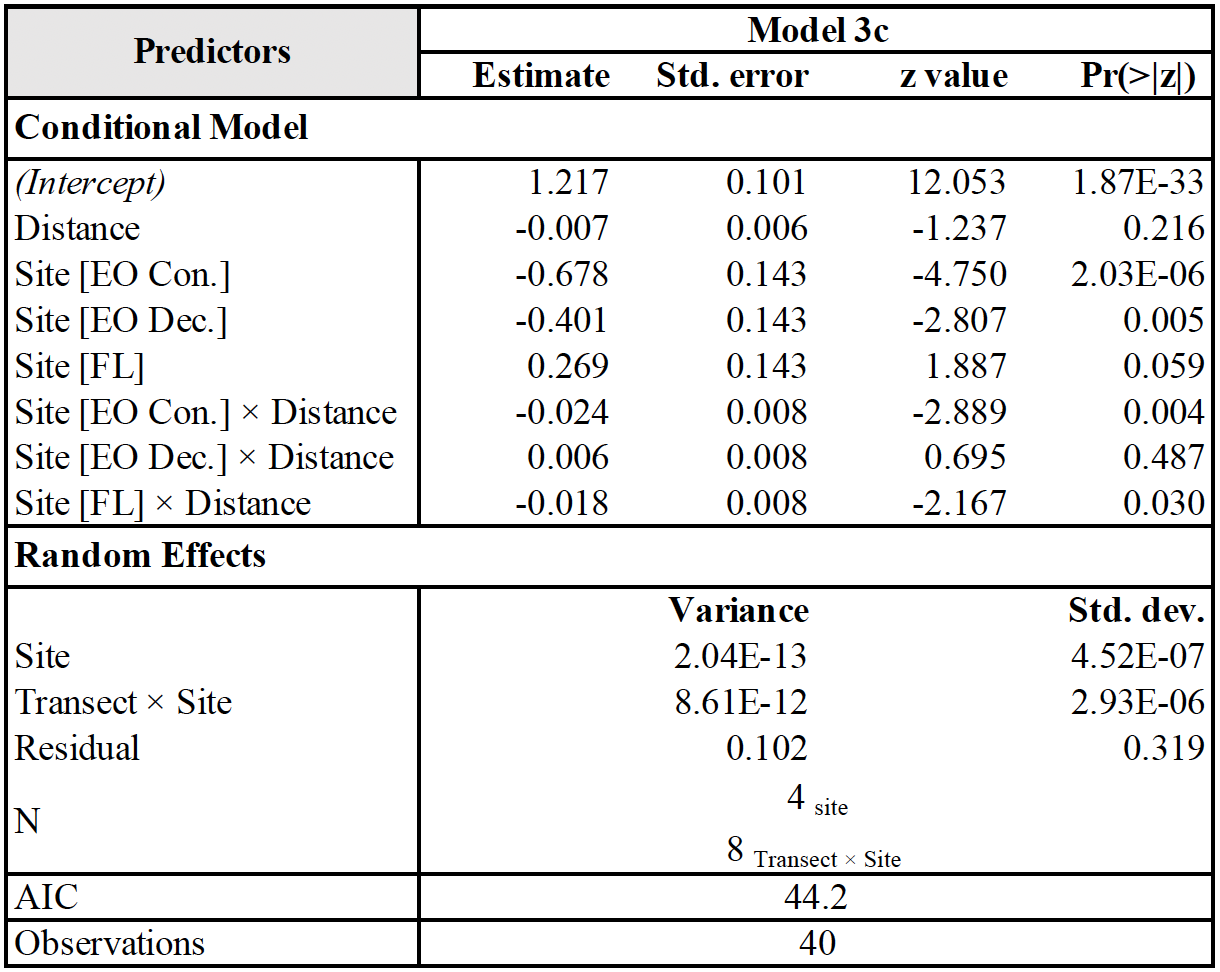

**Table S4.** Results of a generalized linear mixed-effects model analyzing the effects of the site and distance to the field-forest edge on vegetation leaf area index (LAI). Site and transect nested within site were used as random effects. EO = Elginfield Observatory (Con. = Coniferous; Dec. = Deciduous); FL = FRAM Lands. The intercept is the LAI at Baldwin Flats’ edge.

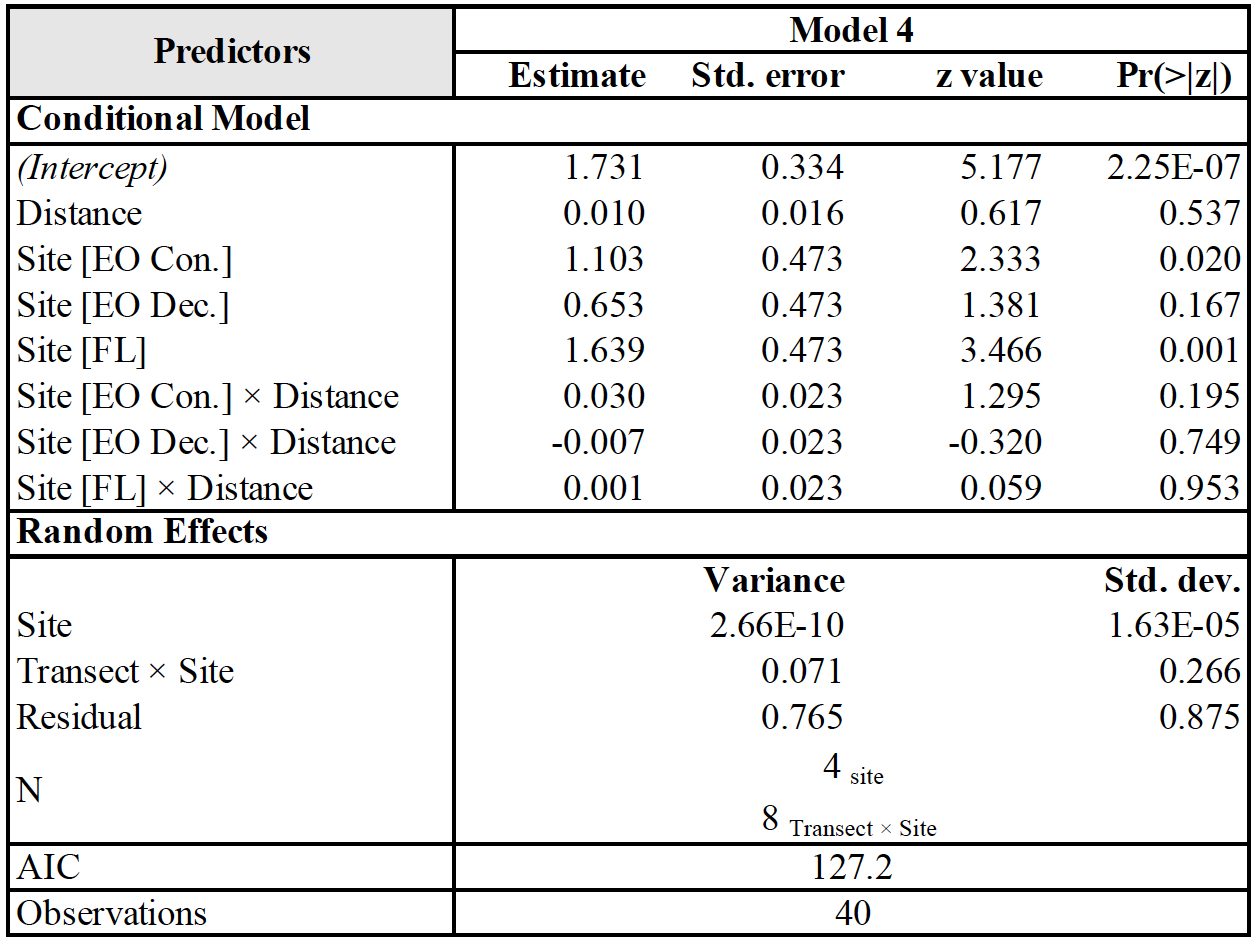

**Table S5a and 5b.** Results of a generalized linear mixed-effects model analyzing the effects of distance to the field-forest edge on (a) macrofauna abundance, and (b) macrofauna richness. Site and transect nested within site were used as random effects. EO = Elginfield Observatory (Con. = Coniferous; Dec. = Deciduous); FL = FRAM Lands. The conditional model intercept is (a) estimated abundance or (b) estimated richness at the edge.

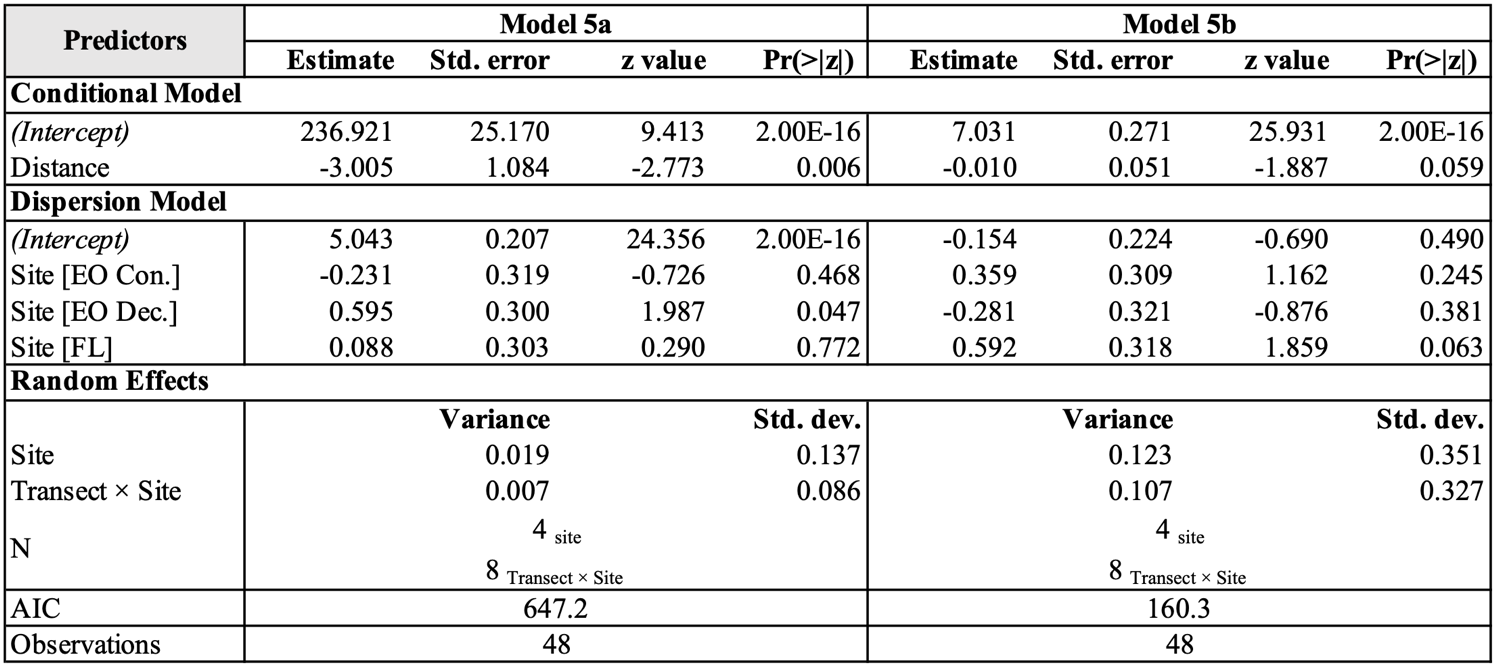

**Table S5c.** Results of a generalized linear mixed-effects model analyzing the effects of distance to the field-forest edge on isopod abundance. Site and transect nested within site were used as random effects. EO = Elginfield Observatory (Con. = Coniferous; Dec. = Deciduous); FL = FRAM Lands. The conditional model intercept is estimated isopod abundance at the edge.

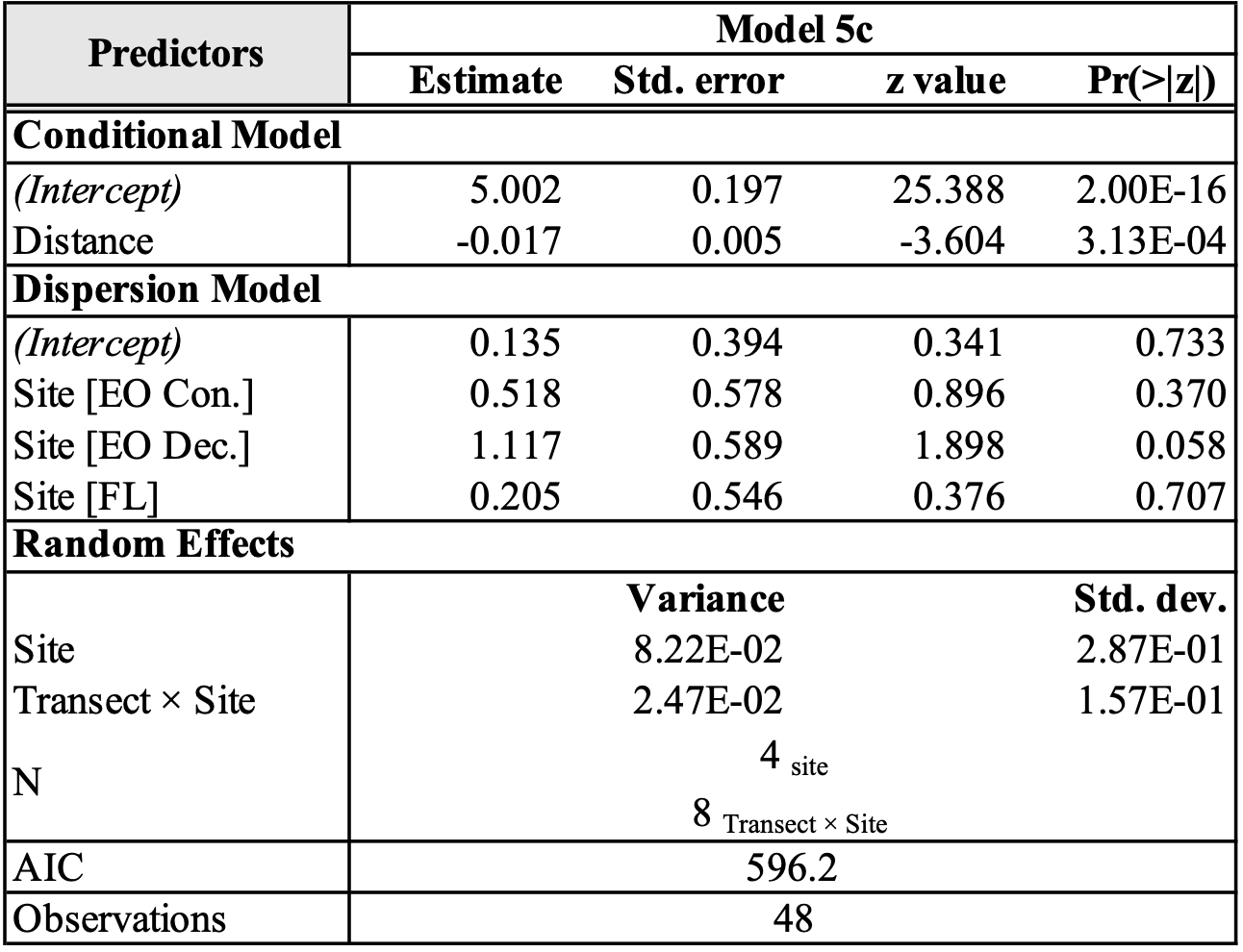

**Table S5d.** Results of a generalized linear mixed-effects model analyzing the effects of distance to the field-forest edge on snail abundance. Site and transect nested within site were used as random effects. EO = Elginfield Observatory (Con. = Coniferous; Dec. = Deciduous); FL = FRAM Lands. The conditional model intercept is estimated snail abundance at the edge.

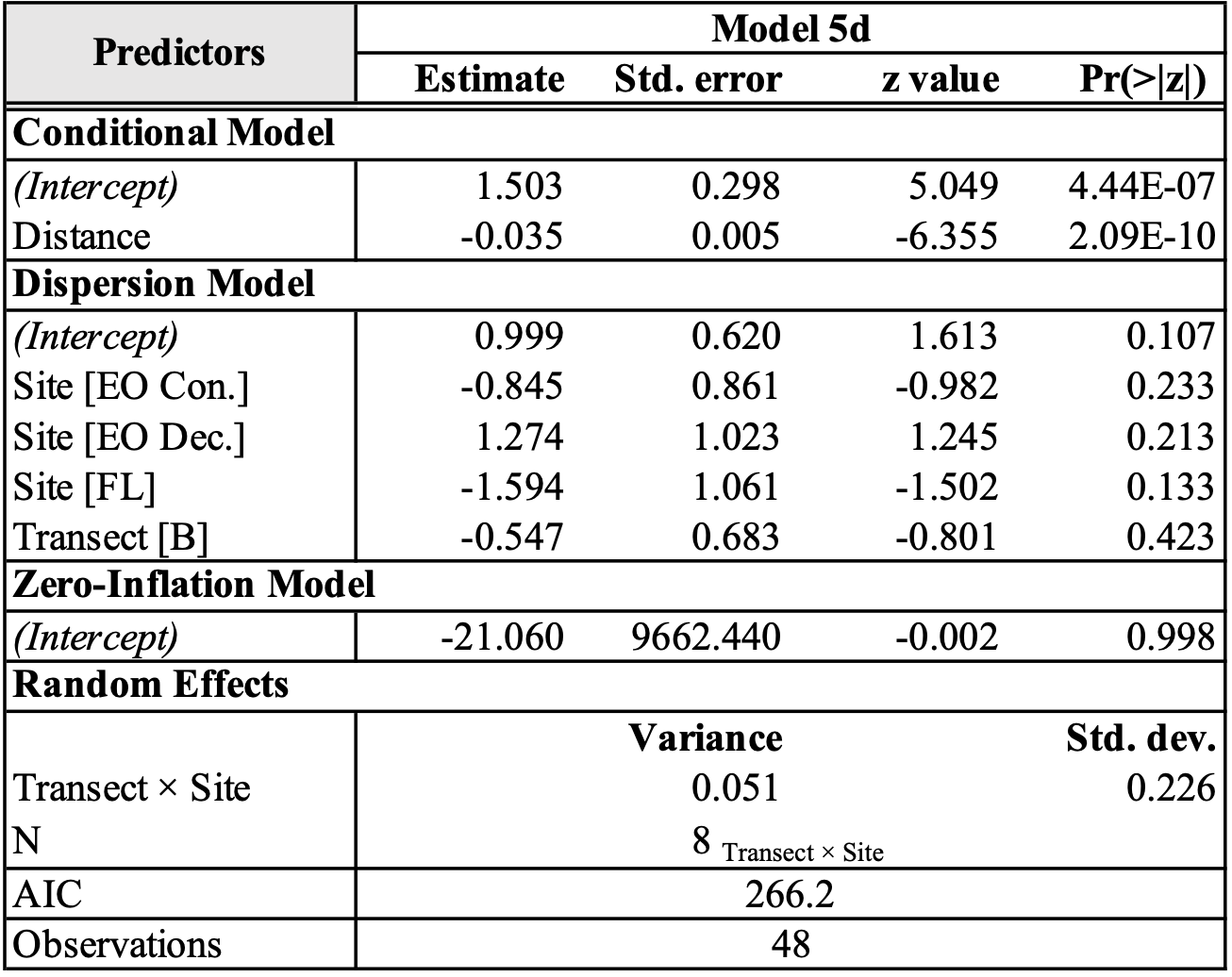

**Table S6.** Results of a generalized linear mixed-effects model analyzing the effects of the interaction of site and distance to the field-forest edge on soil organic matter (SOM). Transect was used as the random effect. EO = Elginfield Observatory (Con. = Coniferous; Dec. = Deciduous); FL = FRAM Lands. The conditional model intercept is estimated SOM at Baldwin Flats’ edge.

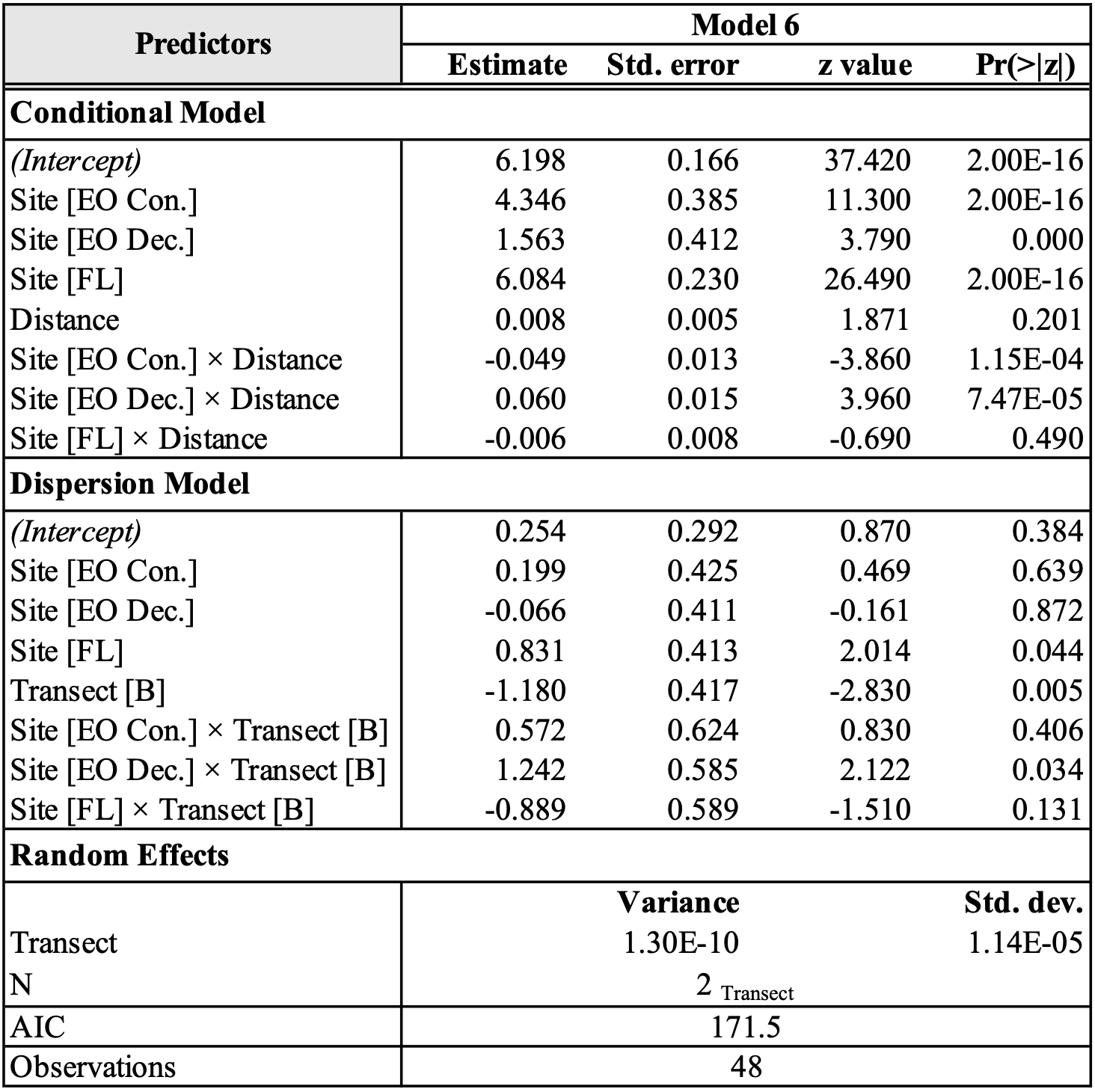
